# Status and prospects of marine NIS detection and monitoring through (e)DNA metabarcoding

**DOI:** 10.1101/2020.05.25.114280

**Authors:** Sofia Duarte, Pedro E. Vieira, Ana S. Lavrador, Filipe O. Costa

## Abstract

In coastal ecosystems, non-indigenous species (NIS) are recognized as a major threat to biodiversity, ecosystem functioning and socio-economic activities. Here we present a systematic review on the use of metabarcoding for NIS surveillance in marine and coastal ecosystems, through the analysis of 42 publications. Metabarcoding has been mainly applied to environmental DNA (eDNA) from water samples, but also to DNA extracted from bulk organismal samples. DNA extraction kits have been widely used and the 18S rRNA and the COI genes the most employed markers, but less than half of the studies targeted more than one marker loci. The Illumina MiSeq platform has been used in >50% of the publications. Current weaknesses include potential occurrence of false negatives due to the primer-biased or faulty DNA amplification and the incompleteness of reference libraries. This is particularly concerning in the case of NIS surveillance, where proficiency in species level detection is critical. Until these weaknesses are resolved, ideally NIS metabarcoding should be supported by complementary approaches, such as morphological analysis or more targeted molecular approaches (e.g. qPCR, ddPCR). Even so, metabarcoding has already proved to be a highly sensitive tool to detect small organisms or undifferentiated life stages across a wide taxonomic range. In addition, it also seems to be very effective in ballast water management and to improve the spatial and temporal sampling frequency of NIS surveillance in marine and coastal ecosystems. Although specific protocols may be required for species-specific NIS detection, for general monitoring it would be vital to settle on a standard protocol able to generate comparable results among surveillance campaigns and regions of the globe, seeking the best approach for detecting the broadest range of species, while minimizing the chances of a false positive or negative detection.

## 1. Introduction

Together with global climate change, overexploitation, pollution and habitat destruction, the spread of non-indigenous species (NIS) is among the major threats to marine and coastal ecosystems (Molnar et al. 2008, Rilov and Crooks 2009). Non-indigenous species, which are introduced from outside of their natural (past or present) distributional range, deliberately or unintentionally by humans or other agents, can spread rapidly in the recipient system, become invasive and displace and out-compete native species (Rilov and Crooks 2009, Simberloff et al. 2013). This can trigger severe ecological changes that threaten ecosystem integrity such as loss of native species and of ecosystem services (Molnar et al. 2008, Rilov and Crooks 2009, Simberloff et al. 2013). One of the major vectors, responsible for the transfer of marine NIS, is transoceanic shipping via ballast waters and hull fouling, or canals connecting large water bodies (Ruiz et al. 2000, Molnar et al. 2008, Hulme 2009). At smaller scales, intraregional or leisure boating may also enhance the spread of NIS (Fletcher et al. 2017, Pochon et al. 2017, Huhn et al. 2020). The ports and marinas, where these vessels dock, often act as hubs for the spread of NIS (Borrell et al. 2017, Grey et al. 2018, Holman et al. 2019). Artificial marine infrastructures (e.g. pontoons), which may be less attractive for native taxa, provide empty niches for opportunistic NIS where they can settle, establish prosperous populations and spread to close vicinities (Dafforn et al. 2015, Olenin et al. 2016).

While visual surveys based on morphological identification of taxa have greatly contributed to ascertain the current status of NIS in marine and coastal habitats (e.g. Borrell et al. 2018, von Ammon et al. 2018b), they require expertise, and are laborious and time consuming. The lack of available expertise on many different taxonomic groups can also pose a major constraint to bioinvasion assessments (e.g. protists: Pagenkopp Lohan et al. 2016, 2017). Moreover, an accurate identification in aquatic systems may be hindered by low visibility waters, as well as by the presence of various life stages not amenable to morphological identification (e.g. eggs, propagules, planktonic larvae, juveniles), because organisms are not large and distinctive (e.g. microalgae, zooplankton, protists) (Pochon et al. 2013, Zaiko et al. 2015b, Pagenkopp Lohan et al. 2016, 2017) or due to low density (e.g. newly arriving NIS) (Darling and Blum 2007).

The detection of an invasive species soon after its introduction, when the population is still confined to a small area and at a low density, maximises the probability of eradication or effective local management (Simberloff 2001, Anderson 2005, Olenin et al. 2011, Pochon et al. 2013). Non-indigenous species are often extremely difficult and expensive to manage, once established (Thresher and Kuris 2004, Olenin et al. 2011). For instance, the EU has spent around 12.5 billion euros annually to control and mitigate the damage caused by NIS (Kettunen et al. 2009). In the U.S., the annual costs of bioinvasions have been estimated at over 100 billion dollars (Pimentel et al. 2001).

Due to the above-mentioned reasons, the development of novel detection methods capable of overcoming some of these challenges becomes a priority in order to contain and eradicate NIS, before they harm ecosystems and result in considerable economic costs (Darling and Blum 2007, Darling and Mahon 2011, Xiong et al. 2016, Darling and Frederick 2018). One such method - DNA metabarcoding - consists on the combination of DNA barcoding (Hebert et al. 2003) with high-throughput sequencing (HTS) and has the potential to bolster current biodiversity assessment techniques (Hajibabaei 2012, Cristescu 2014). In (e)DNA metabarcoding, DNA is extracted from bulk collections of organisms or environmental samples (e.g. water, sediments); one short barcode locus is amplified, and resultant amplicon libraries are sequenced in HTS platforms. Thousands to millions of sequences can be generated in the same reaction, which are processed through a bioinformatics pipeline, where the sequences can be clustered into operational taxonomic units (OTUs) and these OTUs or the sequences directly (exact sequence variants - ESV) are then compared to a reference sequence database to get taxonomic identities, ideally at species level (Hajibabaei 2012, Taberlet et al. 2012, Cristescu 2014, Creer et al. 2016). DNA metabarcoding is rapidly becoming an important approach for direct measurement of biodiversity from environments such as soil, water and air (Baird and Hajibabaei 2012, Taberlet et al. 2012, Deiner et al. 2017) and promises a number of potential benefits over traditional methods. In addition, this approach is easy to implement, which makes DNA metabarcoding one of the tools of choice for the 21st century’s next-generation biodiversity monitoring (Leese et al. 2018, Zinger et al. 2019). The potential for a high sensitivity, greater throughput and accuracy, propensity for standardisation and automation, faster responses and cost effectiveness (Ji et al. 2013, Pochon et al. 2015, Leese et al. 2016, Zinger et al. 2019), makes DNA metabarcoding a particularly well-suited approach for the early detection of NIS and to improve NIS monitoring in biosurveillance programmes (Zaiko et al. 2015a,b, 2016, von Ammon et al. 2018b, Rey et al. 2019, 2020).

Despite the great potential of DNA metabarcoding to improve NIS monitoring in marine and coastal ecosystems (Pochon et al. 2013, Ardura et al. 2015, Zaiko et al. 2015a,b,c, Borrell et al. 2018, Westfall et al. 2020), the widespread and routine implementation of this method is still highly dependent on reliable and cost-effective standardization (Leese et al. 2016, Rey et al. 2020). In this study, we carried out a comprehensive compilation and evaluation of methodologies employed across the full workflow (e.g. sampling design, DNA extraction, targeted genetic markers, sequencing platform, sequence data processing) of DNA metabarcoding applied to eukaryotic NIS detection. We evaluate the current potential for the application of metabarcoding in bioinvasion ecology and assess major weaknesses, challenges and hindrances that still need to be addressed for wider implementation.

## 2. Methodology

A search of the scientific literature was conducted on the Web of Science database (28th March 2020). We searched by topic (which includes words in the title, abstract and keywords) by using all possible combinations of terms that can designate non-indigenous species (i.e. non-indigenous species, invasive species, alien species, exotic species, marine pest) and DNA metabarcoding (i.e. metabarcoding, high throughput sequencing and next generation sequencing) plus the term “marine” or “coastal”, with the exception of “marine pest”, which was used only in combination with the metabarcoding terms. The terms “biosecurity”, “biosurveillance” and “biofouling” were also used in combination with the metabarcoding terms plus “marine” or coastal”. The term “ballast water” was also used in combination with the metabarcoding terms. For more details on the combination of terms used, please see Table S1 of the supplementary material.

The information retrieved from each selected publication included (but was not limited to): i) the geographic area, ii) the sampled substrates and organisms targeted; iii) the sampling design (e.g. n° of sites, n° of sample replicates, n° of technical replicates), iv) the DNA extraction protocols, v) the marker loci targeted and primer pairs used, vi) the sequencing platforms and vii) the bioinformatics pipelines employed to analyse the data.

## 3. Results and discussion

Our literature search yielded 121 papers published between 2010 and 2020. The combinations returning the highest number of matches were: “Invasive species*” or “non-indigenous species*” with “*metabarcoding” (n=31 and 18) or “*high throughput sequencing” (n=20 and 14) plus “*marine”, and “ballast water*” and “*metabarcoding” (n=17) or “*high throughput sequencing” (n=17) (Table S1). After careful inspection we found 34 papers that indeed employed metabarcoding for NIS surveillance in marine and coastal ecosystems and that were retained to perform our analysis. We found 8 additional papers, in our personal collections, that were not displayed during our search, but that have relevant information and, therefore, were included in the analysis (Chain et al. 2016, Ardura et al. 2017, Deiner et al. 2018, Grey et al. 2018, Koziol et al. 2018, Lacoursière-Roussel et al. 2018, Leduc et al. 2019, Suarez-Menendez et al. 2020) (Table S2).

The number of studies employing metabarcoding in NIS surveillance have been increasing over the years, at a rate of ca. 6.3 papers/year (Fig. S1). Top countries, where most of the studies have been conducted, include New Zealand, Canada, Spain, USA and Australia (Fig. S2). This is likely because these countries were early to adopt the use of molecular methods to support management decisions (Darling and Mahon 2011, Xiong et al. 2016). New Zealand is one of the worlds’ countries with the highest rate of non-indigenous species introduction in marine waters (Wood et al. 2013). In this country, the cumulative number of species has risen 10% from 2009 to 2015 (Cranfield et al. 1998, Kospartov et al. 2010, Inglis and Seaward 2016), while in North America, the number of invasive species has more than tripled since the beginning of the century (MEA 2005, Leduc et al. 2019).

### 3.1. Sampled locations, substrates and biological targets

Most of the studies have been conducted in ports or marinas or in the close vicinities (e.g. estuaries), since they are hubs of maritime traffic, where vessels from all continents stop for days or months, unintentionally releasing undesired species to the environment (Table S2, Fig. 1A). In these studies, most sampled substrates were sea water and ships ballast water and the majority applied metabarcoding to environmental DNA (eDNA) (Table S2, Fig. 1B) (e.g. Borrell et al. 2017, Deiner et al. 2018, Grey et al. 2018, Rey et al. 2019, Wood et al. 2019, Suarez-Menendez et al. 2020) or to zooplankton (e.g. Zaiko et al. 2015a,b, Abad et al. 2016, Ghabooli et al. 2016, Darling et al. 2018, Lin et al. 2020).

**Figure 1.**
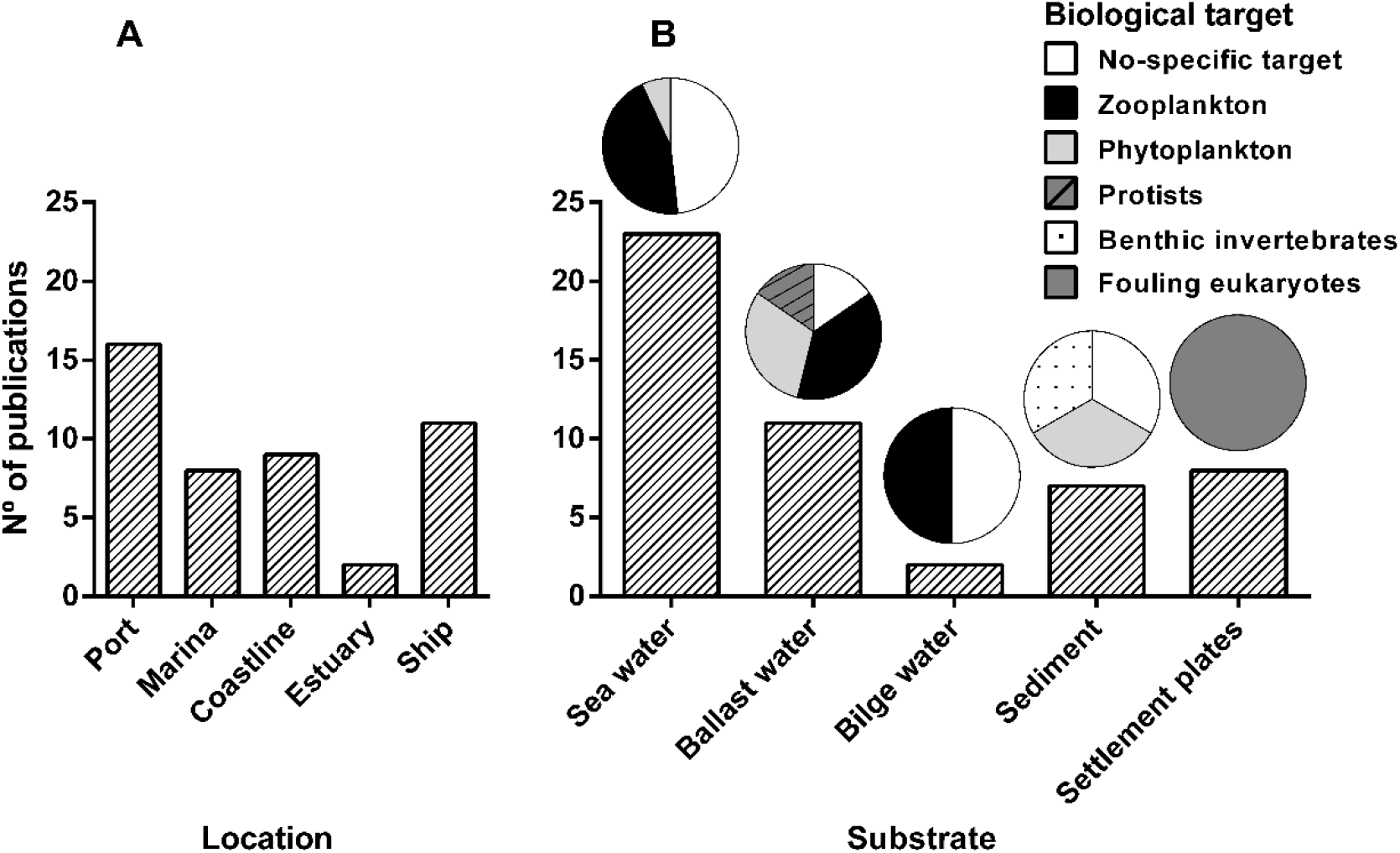
Type of locations (A), sampled substrates and biological target (B) in published studies pertaining the application of DNA metabarcoding in NIS surveillance in marine and coastal ecosystems.

Environmental DNA can be extracted directly from environmental samples, such as water and sediments, and can originate from organisms’ body parts, mucus, tissues, cells, secreted faeces or extracellular DNA, leaking to the environment through the normal life or upon death of organisms (Turner et al. 2014, Sassoubre et al. 2016, Holman et al. 2019). Targeting eDNA can be particularly advantageous to provide a rapid snapshot or baseline information of local species composition and for early detection of NIS, without the need for sampling individual organisms (Ficetola et al. 2008, 2018, Hajibabaei et al. 2011, Taberlet et al. 2012, Singer et al. 2019). Environmental DNA has been the exclusive biological target in 11 biosurveillance studies employing metabarcoding (Table S2). Repeated eDNA surveys may have also the potential to be used to evaluate long-term biodiversity changes and NIS range expansions (Lacoursière-Roussel et al. 2018, Leduc et al. 2019), or to monitor NIS in ballast waters or to test the efficiency of treatment systems (Ardura et al. 2015, Rey et al. 2019, Lin et al. 2020). However, studies employing both eDNA and bulk organismal samples suggest that eDNA alone cannot replace organismal sampling (e.g. Leduc et al. 2019, Huhn et al. 2020, Rey et al. 2020, Westfall et al. 2020). Currently, it would probably work best as an early alert system or exploratory method to be used as a complement to conventional approaches such as *in situ* visual confirmation (Borrell et al. 2018, Holman et al. 2019, Rey et al. 2020), specimen collection (Leduc et al. 2019, Huhn et al. 2020) or followed by active surveillance with digital drop PCR (ddPCR) or quantitative PCR (qPCR) (Wood et al. 2019).

Zooplankton has been by far the most targeted organisms in marine NIS surveillance using metabarcoding (Fig. 1B). Zooplankton sampling has been conducted in ships ballast water (Zaiko et al. 2015a,b, Ghabooli et al. 2016, Darling et al. 2018, Lin et al. 2020), boats bilge water (Fletcher et al. 2017), and in sea water in ports and marinas or nearby natural habitats (Pochon et al. 2013, Zaiko et al. 2015c, Abad et al. 2016, Brown et al. 2016, Chain et al. 2016, Stefanni et al. 2018, Couton et al. 2019, Leduc et al. 2019, Pagenkopp Lohan et al. 2019, Rey et al. 2020, Westfall et al. 2020). The monitoring of larvae in plankton samples in ports or in the close vicinities, may provide key information about NIS introduction status or detect their presence at an earlier stage (Couton et al. 2019). However, targeting dispersive life stages of NIS in the water column has been mostly impractical in routine monitoring using traditional methods. The small size and cryptic nature of planktonic taxa and larval stages, as well as the lack of specific taxonomic expertise, makes zooplanktonic organisms well-suited to be identified through metabarcoding. In the pioneer study of Pochon et al. (2013), and despite the co-occurrence of a large number of other eukaryotes, a single spiked *Asterias amurensis* larvae was detected in plankton samples through metabarcoding. Several other planktonic larval NIS stages have been exclusively detected through metabarcoding (Zaiko et al. 2015c, Fletcher et al. 2017, Couton et al. 2019), including short-lived larval stages (e.g. *Bugula neritina*, *Corella eumyota*; Couton et al. 2019), which otherwise may have remained unnoticed for long time. Metabarcoding is also becoming an efficient tool for the detection and identification of many phytoplankton species that do not fall into the regulated 10-50 μm size class (Zaiko et al. 2015a,b, Wright et al. 2019), and in the identification of parasitic protists, particularly toxic dinoflagellates, transported in the ballast water of ships (Pagenkopp Lohan et al. 2016, 2017, Petri et al. 2019, Shang et al. 2019). Metabarcoding can also greatly benefit the identification of hard-substrate dwelling organisms, that become part of biofouling communities in natural and artificial substrates (e.g. settlement plates, Table S2, Fig. 1B) (Pochon et al. 2015, Zaiko et al. 2016, Koziol et al. 2018, von Ammon et al. 2018a,b, 2019, Wood et al. 2019, Rey et al. 2020). For instance, many metazoan genera containing NIS or cryptogenic species were detected through metabarcoding from scarce genetic material in early fouling communities, in settlement plates deployed in New Zealand’s marinas (e.g. *Ciona* sp., Pochon et al. 2015; *Watersipora* sp., *Ciona* sp., *Molgula* sp. and other Bothryllid ascidians, Zaiko et al. 2016). In addition, DNA-based results were consistent with the traditional approach for higher taxonomic ranks, but DNA metabarcoding was able to detect many more taxa at the genera/species level (Zaiko et al. 2016). This is extremely relevant for the early detection of NIS in vessels or equipment from high-risk regions that can be rapidly checked prior to arrival at destination, when the species are still at low densities and morphologically unidentifiable (Pochon et al. 2015).

Compared to water, the collection of sediments to survey eukaryotic organisms has received much less attention, and to our knowledge has been attempted only in 6 studies (Table S2, Fig. 1B, Koziol et al. 2018, Holman et al. 2019, Leduc et al. 2019, Shang et al. 2019, Shaw et al. 2019, Rey et al. 2020). Metabarcoding of eDNA extracted from sediments can provide a rapid screen for a wide range of taxa in ballast tanks and marine port sediments (Holman et al. 2019, Shang et al. 2019, Shaw et al. 2019). In particular, cysts of toxic dinoflagellates and diatoms resting stages are prime candidates for successful transport in sediments from ballast tanks (McCarthy and Crowder 2000, Shang et al. 2019). Early detection through metabarcoding can help prevent spread to new locations, where they can result in toxic blooms and have devastating impacts on ecosystems (Shang et al. 2019, Shaw et al. 2019). On the other hand, benthic invertebrates have been only surveyed in two studies using metabarcoding in NIS surveillance (Leduc et al. 2019, Rey et al. 2020). Benthic habitats can be very complex and benthic communities very patchy, which increases the possibility of missing taxa when sampling invertebrates from sediments (Leduc et al. 2019). In addition, many marine benthic NIS have a biphasic bentho-pelagic life cycle and targeting their pelagic larval stages in zooplankton its easier and may allow early detection, before establishment and potential spread (Zaiko et al. 2015c, Couton et al. 2019).

### 3.2. Sampling design

We found that most of the studies have sampled multiple ports or marinas (e.g. Brown et al. 2016, Chain et al. 2016, Borrell et al. 2017, Leduct et al. 2019), or estuaries (Borrell et al. 2018), and multiple sites within each port/marina/estuary (Abad et al. 2016, Brown et al. 2016, Leduc et al. 2019, Rey et al. 2020). In order to detect the majority of taxa in a given location, in particular rare and/or patchy species, biodiversity surveys require comprehensive spatial and temporal sampling and screening of different types of substrates (e.g. water, sediment, bulk samples; Koziol et al. 2018, Huhn et al. 2020, Rey et al. 2020, Westfall et al. 2020). However, the sampling design varied significantly across studies and very few have compared the effects of the sampling effort or the use of different substrates in NIS detection (see Zaiko et al. 2016, Koziol et al. 2018, Lacoursière-Roussel et al. 2018, Wood et al. 2019, Rey et al. 2020, Westfall et al. 2020) (Table S2). Conducting trial experiments would be particularly helpful to assess the appropriateness of the sampling strategies employed and how these factors can affect NIS detection at particular sites in coastal and marine ecosystems (Deiner et al. 2018, Koziol et al. 2018, Pagenkopp Lohan et al. 2019, Huhn et al. 2020, Rey et al. 2020). For instance, based on an extensive survey conducted in the port of Bilbao (Spain), Rey et al. (2020) found a strong influence of the sampling site in the recovered biodiversity, suggesting that spatially comprehensive sampling was crucial to recover biodiversity, and to increase the success of NIS detection. Also, higher sampling frequency has been shown to increase the diversity recovered, as well as the ability to detect NIS (Chain et al. 2016, Lacoursière-Roussel et al. 2018, Couton et al. 2019, Rey et al. 2020, Westfall et al. 2020). Because marine communities may experience considerable shifts in composition across seasons and reproductive periods (Sutherland and Karlson 1977, Bachelet et al. 2009), narrow spatial and temporal sampling scales are recommended. However, finer sampling scales will also be reflected in greater associated costs. In the case of NIS detection, a compromise may be found through the selection of specific seasons as for example, shortening sampling to spring and late summer in temperate latitudes, as suggested by Rey et al. (2020). To allow such sampling optimization, the metabarcoding approach would still need to be tested across a wide geographic scope in the survey of a broad variety of NIS present in natural communities of different latitudes and environmental settings (Table S2, Fig. S2).

In our review, we found that DNA metabarcoding has been applied to DNA extracted from bulk organismal samples or to eDNA, namely water (see previous sub-section). However, marine NIS can have different life history traits, such as habitat preferences and life cycles, which will likely influence the detection of taxa in a given substrate. Preliminary studies that have compared different substrates found that the total diversity recovered, including distinct subsets of NIS within specific taxonomic groups, have been accomplished only when combining different methods (Koziol et al. 2018, Leduc et al. 2019, Huhn et al. 2020, Rey et al. 2020, Westfall et al. 2020). For example, Koziol et al. (2018) found that fish were preferentially detected in water, nematodes in sediments and crustacea in planktonic tows, in two Australian coastal ports. Bryozoans and ascidians are encrusting organisms and thus highest detection rates are expected to be found in settlement plates or close to hard substrata. Rey et al. (2020) also found distinct communities surveyed in the port of Bilbao (Spain) by using different sampling methods (i.e. PVC plates deployment, plankton nets, water and sediment sampling). Combining different sampling methods may be crucial to maximize NIS detection at a particular site (Koziol et al. 2018, Huhn et al. 2020, Rey et al. 2020, Westfall et al. 2020). Within each site, the use of several replicates per substrate may also be better suited to maximize the chances of NIS detection (Koziol et al. 2018, Holman et al. 2019, Westfall et al. 2020).

### 3.3. Sample pre-processing before DNA extraction

We found very few studies that have analysed the effects of sample pre-processing in marine NIS surveillance studies through DNA metabarcoding (but see Deiner et al. 2018, Pagenkopp Lohan et al. 2019) (Table S2, S3). Yet, according to some studies, processing steps prior to DNA extraction, such as refrigeration or freezing, homogenization or choice of filters used in water filtration, may strongly influence NIS detection efficiency (Deiner et al. 2018, von Ammon et al. 2018b, Pagenkopp Lohan et al. 2019, Shaw et al. 2019). For example, Deiner et al. (2018) mixed 13 L of sea water collected in the port of Southampton (UK) in a container, that was maintained under continuous stirring before water filtration. Von Ammon et al. (2018b) used a stomacher to macerate biofouling organisms growing on PVC plates. Also, Shaw et al. (2019) homogenised all sediment samples collected from ballast water tanks from ships arriving to Australian ports, prior to DNA extraction. These procedures ensure homogeneity among samples and may increase the recovered diversity, thereby improving the chances of early NIS detection. For instance, in a survey of zooplankton in USA bays, Pagenkopp Lohan et al. (2019) found that homogenizing samples with a mortar and a pestle, prior to DNA extraction, greatly increased the detection of rare or low abundance OTUs.

Different water volumes and types of filters (with different composition and pore size) have also been used to analyse eDNA from water (Table S3, Fig. 2). Since marine water masses are enormous and highly dynamic (exposed to tides, currents), the eDNA of rare species may be highly diluted. Thus, sampling a large volume of water may be needed to increase the chances of species detection, but this can constitute a practical problem in routine surveys (Borrell et al. 2017). A large volume of water cannot be easily refrigerated or frozen until analysis, so it should be filtered *in situ* (Jerde et al. 2011, Goldberg et al. 2016, Hinlo et al. 2017, Majaneva et al. 2018). Another option could be chemical preservation with cationic surfactants (e.g. benzalkonium chloride), which can protect eDNA in water samples with less effort and equipment (Yamanaka et al. 2017). However, most authors opted to filter smaller volumes (30 mL to 6 L) (Table S3), choosing a compromise between feasibility, in terms of sample storage and processing time, and taxa representativeness for a given location. One possibility to increase the throughput volume of capture filters and reduce filtration time may be provided by the use of pre-filters with a larger pore size, followed by a second filtration of a smaller volume through a smaller pore (Ardura et al. 2015, von Ammon et al. 2019, Wood et al. 2019) (Table S3). Filtering small volumes through independent filters, to avoid filter clogging, and pooling results together during sequence analysis, can be another valid option (Leduc et al. 2019, Suarez-Menendez et al. 2020).

**Figure 2.**
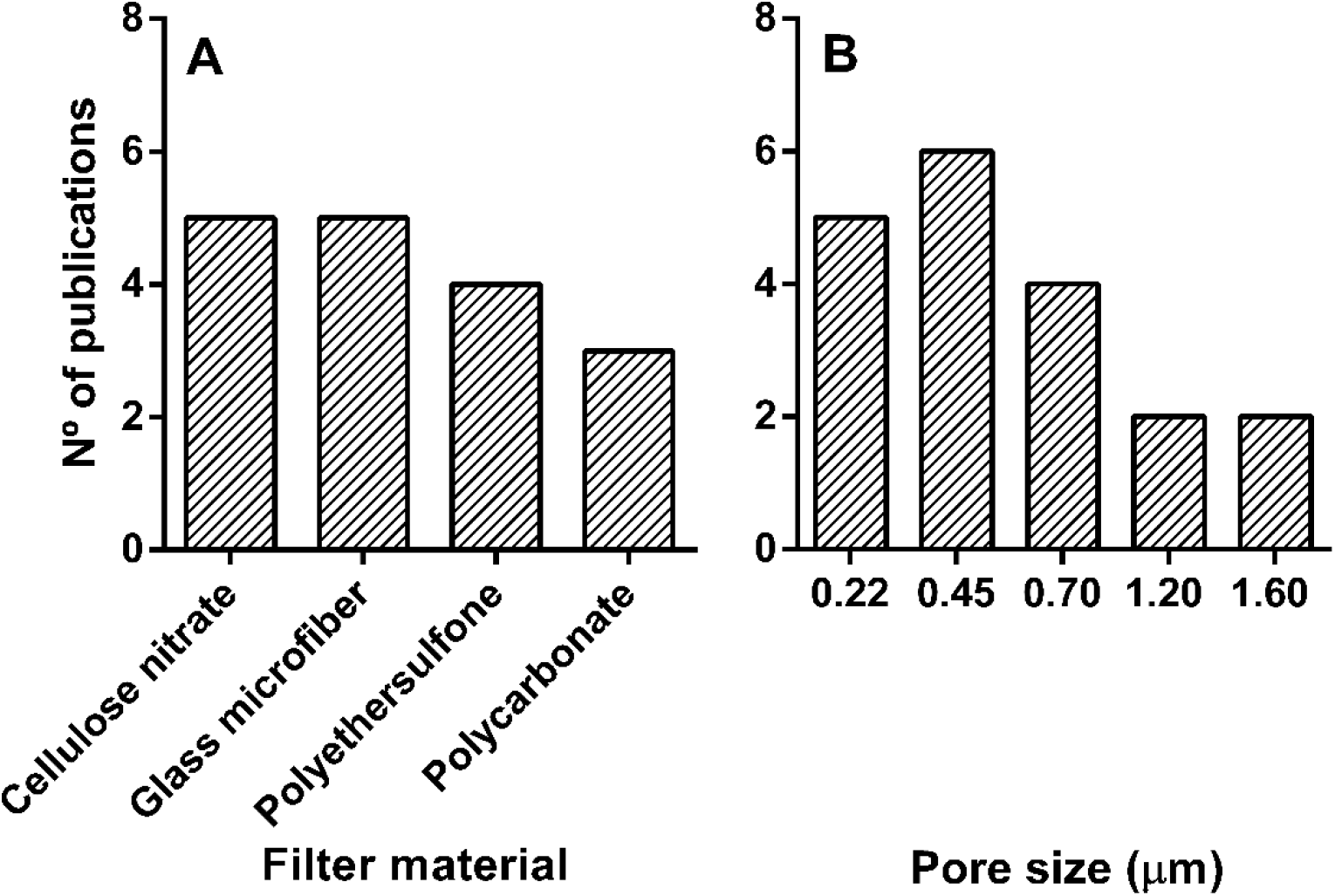
Filter type (A) and pore size (B) used to filter water samples in published studies pertaining the application of DNA metabarcoding in NIS surveillance in marine and coastal ecosystems.

The depth at which the water is sampled is another technical aspect that varied across studies (Table S3). Most studies sampled surface water (10-30 cm) (Borrell et al. 2017, 2018, Deiner et al. 2018, Grey et al. 2018, Koziol et al. 2018, Holman et al. 2019, Huhn et al. 2020, Suarez-Menendez et al. 2020), but many others sampled different depths to increase representativeness (Lacoursière-Roussel 2018, Leduc et al. 2019, von Ammon et al. 2019, Rey et al. 2020, Westfall et al. 2020). Vertical eDNA distribution in the water column may vary as a function of several factors (e.g. life cycle of species, transport, haloclines, and other complex hydrodynamic processes) (Jeunen et al. 2019) and DNA may be less degraded at greater depths. Thus, ideally, water column stratification and vertical community structuring should be taken into account in eDNA metabarcoding surveillance studies.

The amount of sediment used for eDNA extraction is another factor that varied considerably among studies (Table S4, 0.25-10 g), depending on the target organisms (Koziol et al. 2018, Holman et al. 2019, Shang et al. 2019, Shaw et al. 2019). In general, studies targeting macro-eukaryotes used higher amounts of sediment (5-10 g) (Pochon et al. 2013, Holman et al. 2019), while lower amounts have been used for targeting micro-eukaryotes (0.25-0.45 g) (Shang et al. 2019, Shaw et al. 2019).

Concerning the effects of the filter material and pore size, to our knowledge there was only one study that tested its effects on NIS eDNA detection in water samples (Deiner et al. 2018). The authors found that the filter material (glass microfiber *versus* cellulose nitrate), but not the pore size (0.2 to 1.2 μm), affected OTUs richness recovered using eDNA metabarcoding from seawater in the port of Southampton (UK) (Deiner et al. 2018). Both types of filters and a 0.45 μm pore size, have been widely used in NIS surveillance through eDNA metabarcoding (Fig. 2), but a greater OTU recovery was found with cellulose nitrate filters in the study of Deiner et al. (2018). On the other hand, increasing the pore size (e.g. 0.8 μm) may greatly improve filtration time without compromising the taxa detected from eDNA (Li et al. 2018). Thus, more targeted studies addressing technical details regarding eDNA sampling collection and processing are required, in particular the choice of membrane filters, where both the material and pore size can play important roles in eDNA capture (Deiner et al. 2018). Handling and exposure to outside stress may be reduced by using enclosed filters, that may also be better suited to perform on-site filtration, contributing to more accurate eDNA results (Spens et al. 2017, Li et al. 2018). Yet, to our best knowledge, they were employed only in two studies using eDNA metabarcoding in NIS surveillance (Holman et al. 2019, Rey et al. 2020).

### 3.4. DNA extraction

Concerning DNA extraction, most of the studies have used DNA isolation kits (39 publications, Table S2, Fig. 3). In particular, MoBio kits (further acquired by Qiagen) have been used in 32 out of 42 studies (Table S2, Fig. 3), which have been reported to yield high quality DNA for barcoding or metabarcoding applications (Borrell et al. 2017, Djurhuus et al. 2017). In addition, PowerSoil and PowerMax soil DNA isolation kits are particularly useful for sediment samples and other complex samples due to the removal of inhibitors, that can affect considerably the amount of diversity detected (Hermans et al. 2018). Commercial DNA isolation kits have the advantage of requiring limited and smaller amounts of chemicals, shortened isolation steps, and achieving results quicker, but, usually, have high costs associated. Cost-effective extraction methods may increase the ability to process a large number of samples and can also produce good results, but may need extra purification steps (Deiner et al. 2018, Lacoursière-Roussel et al. 2018, Leduc et al. 2019).

**Figure 3.**
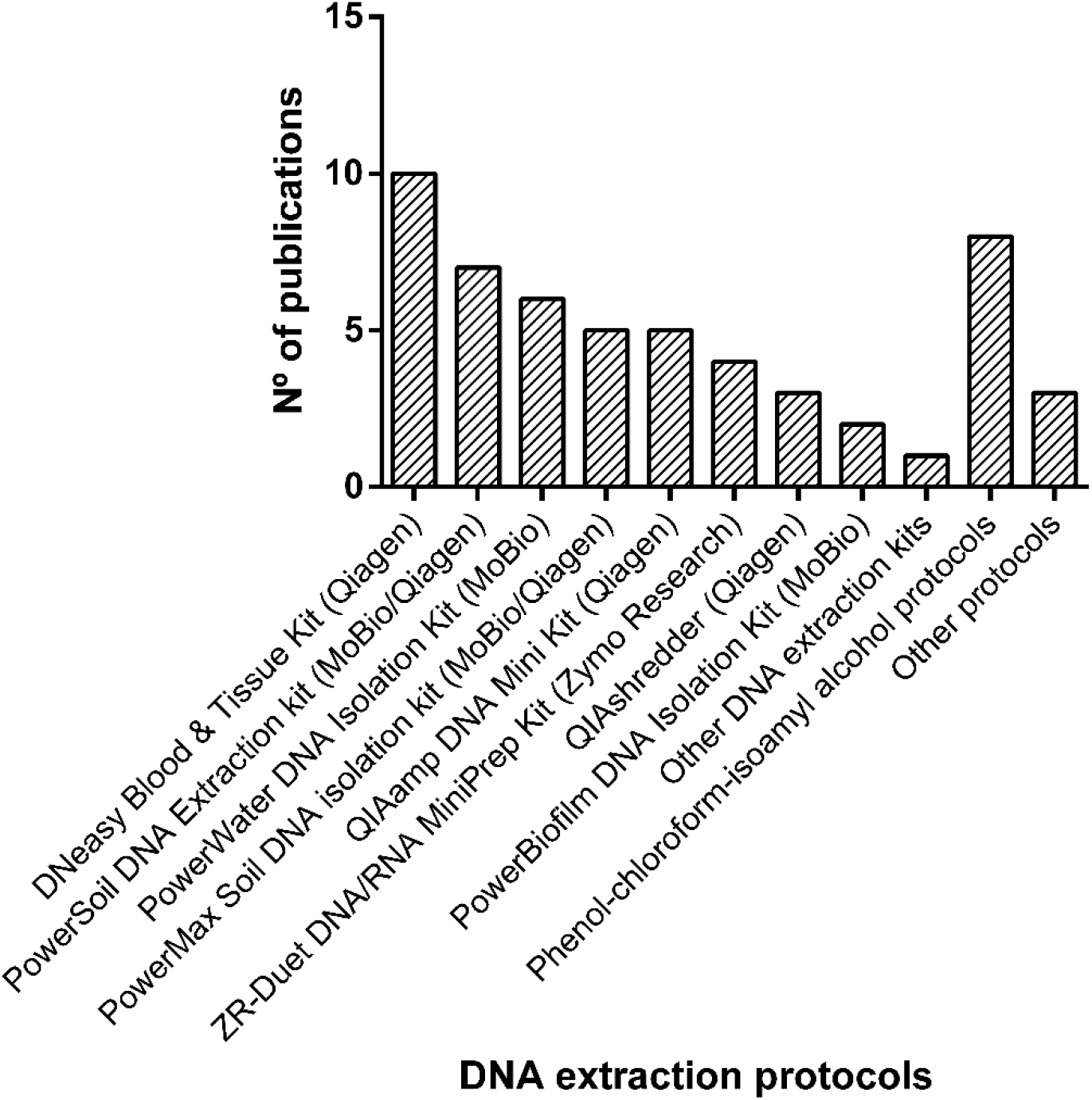
DNA extraction protocols used in published studies pertaining the application of DNA metabarcoding in NIS surveillance in marine and coastal ecosystems.

To our best knowledge only one study assessed the efficiency of different DNA extraction methods in NIS detection rates (Deiner et al. 2018). Deiner et al. (2018) found that both filter material (see the previous sub-section) and DNA extraction protocols significantly affected the detection rate of the invasive ascidian *Styela clava*, in the port of Southampton (UK). Many organisms may indeed remain unidentified using DNA metabarcoding due to deficient DNA extraction (e.g. those that are rare, small or calcified). The failure to detect through DNA metabarcoding the polychaete worm *Spirorbis* sp. on PVC plates deployed in a New Zealand’s port, was attributed to inefficient DNA extraction, though faulty amplification cannot also be excluded (Zaiko et al. 2016).

The inclusion of technical replicates, of the same sample, during DNA extraction and for the same DNA extract during PCR (e.g. Brown et al. 2016, Chain et al. 2016, Ghabooli et al. 2016, Pagenkopp Lohan et al. 2016, Lanzén et al. 2017, Koziol et al. 2018, Lacoursière-Roussel et al. 2018, Shaw et al. 2019, Rey et al. 2020) should also be considered to disentangle the effects of technical variance and increase the rate of NIS detection, particularly if DNA concentrations are low. For instance, Pagenkopp Lohan et al. (2019) found a significant effect on the recovered zooplankton diversity, collected at San Diego bays (USA), by increasing the number of DNA extraction replicates. Deiner et al. (2018) used a statistical power analysis prior of their study to determine the number of experimental replicates needed for DNA extraction and to increase the confidence of their results. Since DNA extraction methods can strongly influence the biotic composition of the samples (Hermans et al. 2018), different methods may need standardization at some steps of the purification protocols to yield comparable results. For example, Koziol et al. (2018) used a standardized silica-binding purification process across different DNA extraction methods, to better compare the recovered diversity from different substrates.

### 3.5. Use of controls

The inclusion of negative and positive controls, through the complete DNA metabarcoding workflow (i.e. sampling, DNA extraction, PCR and sequencing), can facilitate the detection of contamination or potential artefacts introduced at any step (Zinger et al. 2019). Negative controls (i.e. DNA storage or extraction buffers, sterilized ultra-pure water) have been frequently used in NIS surveillance studies employing (e)DNA metabarcoding, from field sampling until sequence processing. On the other hand, positive controls (i.e. DNA from mock communities or samples with *a priori* known composition) have not been so frequently used (Table S2). However, positive controls or prior expectations of taxa occurring in the system (i.e. NIS updated lists), are recommended to evaluate the effectiveness of the metabarcoding approach (Zinger et al. 2019). For instance, Zaiko et al. (2016) included DNA samples from 3 NIS (*Sabella spallanzanii*, *Ciona savignyi* and *Perna perna*) as internal quality controls. After purification and quantification of PCR amplicons, these were combined at 3 distinct concentrations and 2 replicates of each were incorporated into the HTS library. The taxonomic assignment of the internal control confirmed the robustness of the sequencing and analytical pipeline used, with more than 99.9% of the sequences being assigned to the target taxa (Zaiko et al. 2016). This procedure has been adopted in posterior studies (Fletcher et al. 2017, Pochon et al. 2017).

PCR inhibition controls may also be applied to ensure that extracted DNA is free of inhibitors, by using qPCR (Holman et al. 2019, Wood et al. 2019, Westfall et al. 2020). Another possibility for implementing PCR inhibition controls consists in the introduction of a known amount of DNA from a non-target species in the DNA template, prior to amplification. The introduced DNA must be from an organism very unlikely to occur in the surveyed environment, or made of artificially synthetized DNA, and known *a priori* to amplify with the primers in use. For instance, Holman et al. (2019) filtered water from a marine tropical aquarium containing a variety of hard and soft corals, molluscs and tropical fish, as a positive control, while minimizing the risk of cross contamination that may occur when using target species. In addition, the authors also added DNA from *Microcosmus squamiger* (a species not currently known to occur in UK waters) to the filter before DNA extraction, to control for inhibitors that may be found in marinas and harbours.

Inhibition can be overcome through the addition of PCR enhancers (e.g. BSA, Suarez-Menendez et al. 2020), through dilution of sample DNA (not always advisable when working with low concentrations) (Wood et al. 2019), or through additional inhibitor removal steps after DNA extraction (Deiner et al. 2018, Wright et al. 2019).

The inclusion of these controls through the analytical chain of the metabarcoding approach are extremely important to better understand the nature of two types of errors: i) the Type-I errors or false positives namely due to contaminations (field, DNA extraction, PCR), “tag/index jumps”, or PCR and sequencing errors, and ii) the Type-II errors or false negatives, due to, for example partial sampling, DNA extraction, amplification or sequencing bias (Zinger et al. 2019). Both types of errors, that may often arise during the use of DNA metabarcoding, can trigger action or inaction when not required, causing a potentially large burden on entities responsible for invasive species mitigation and control (Pochon et al. 2017). The use of the morphological approach or at least coarse visual inspections (e.g. Zaiko et al. 2015b,c, 2016, Abad et al. 2016, Fletcher et al. 2017, Huhn et al. 2020, Borrel et al. 2018, Holman et al. 2019) can be very important to better detect metabarcoding false negatives and to understand its nature (e.g. incompleteness of reference libraries *versus* methodological limitations such as the above-mentioned). However, these have been considered in less than half of the studies (Table S2).

In addition, finding evidence of DNA from a species, particularly in environmental samples, may not necessarily means that it belongs to a living organism (Pochon et al. 2017, von Ammon et al. 2019). Only living organisms can indeed become invaders and pose risks to ecosystems. The use of RNA may offer an alternative for identifying living organisms in samples, since it is directly linked with active gene expression of metabolic pathways and degrades rapidly upon cell death (Pawlowski et al. 2014, Pochon et al. 2017, von Ammon et al. 2019, Wood et al. 2020). In a recent study the authors found eRNA persisting in the marine environment longer than expected, which may provide new opportunities for improving biodiversity surveys through reducing false positives (Wood et al. 2020). However, working with RNA requires particular storage conditions of samples and workflow protocols which are expensive, time-consuming and less frequently used (von Ammon et al. 2019, Wood et al. 2019).

### 3.6. Marker loci and primer pairs

The identification at higher taxonomic levels (e.g. family or order), is often enough to broadly estimate biodiversity through DNA metabarcoding (e.g. Creer et al. 2010, Bik et al. 2012). But, in the case of NIS detection, species-level identification is often mandatory, and highly dependent of the marker and choice of primer pair. The so-called “universal” or “near universal” primers, targeting the nuclear SSU or 18S rRNA (18S) and the mitochondrial cytochrome c oxidase subunit I (COI) genes, have been the most widely used in studies of DNA metabarcoding targeting NIS in marine and coastal ecosystems (in 31 and 22 publications, respectively, Table S2, Fig. 4).

**Figure 4.**
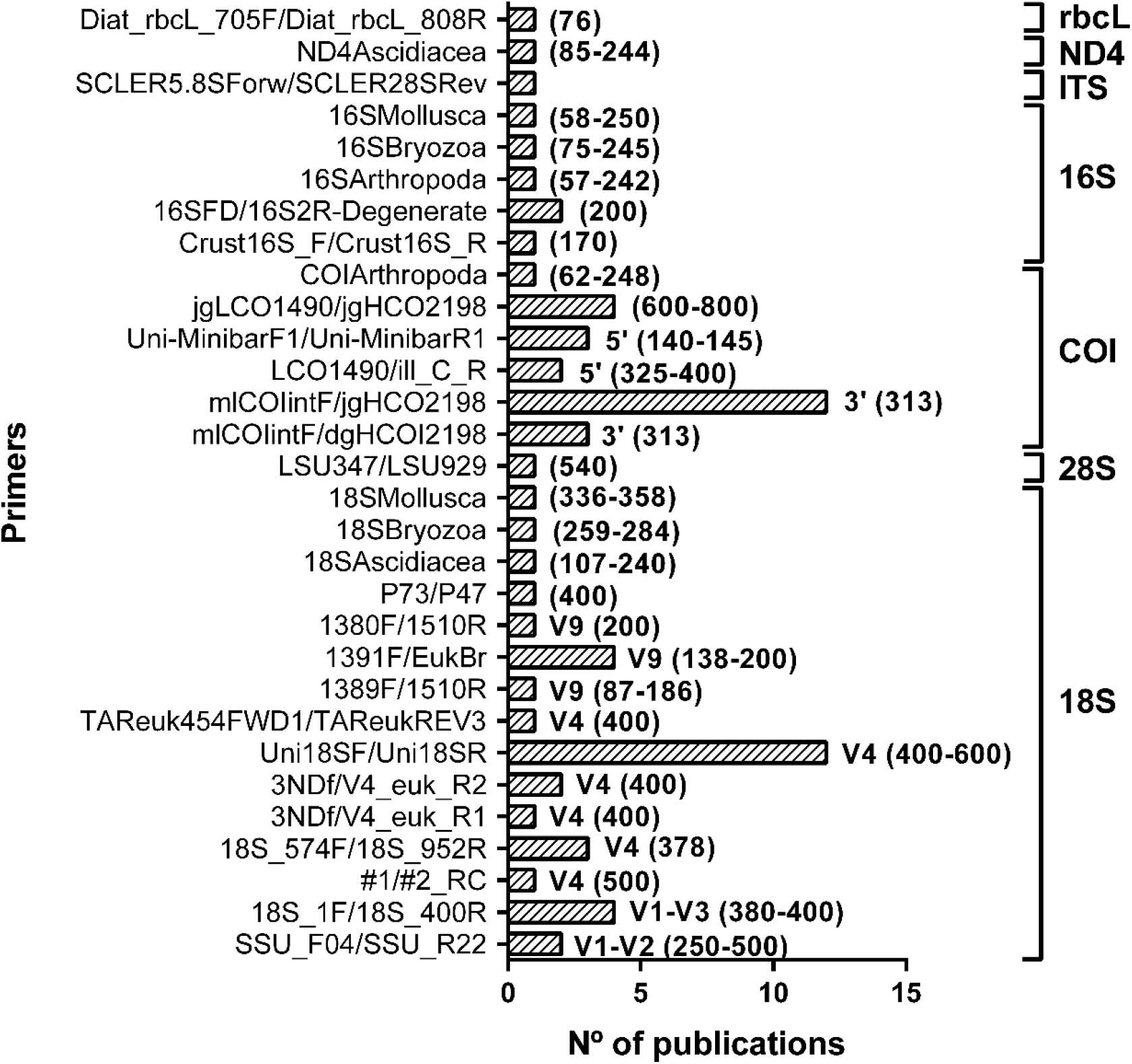
Target loci and primer pairs used in published studies pertaining the application of DNA metabarcoding in NIS surveillance in marine and coastal ecosystems. Within each marker, specific regions used are indicated in the top of each bar, as well as the approximate amplicon length between brackets.

Different regions of 18S have been predominantly used when considering marine microbial eukaryotic diversity (Amaral-Zettler et al. 2009), since current primers allow assessment of both metazoan and protist communities (Darling et al. 2018). The 18S region is also the formal primary marker adopted for barcoding protists (Pawlowski et al. 2012). Most of the studies employing 18S targeted the V4 hypervariable region (Fig. 4), since current primers have high amplification success, covering major invertebrate groups of Crustacea, Mollusca and Tunicata (Zhan et al. 2013). High amplification success was also found for additional relevant taxonomic groups such as Ciliophora, Alveolata, but also for main groups of algae, i.e. Chlorophyta or Rhodophyta (Briand et al. 2018). Although better successful amplification rates are usually achieved with 18S due to higher conservation of primer-binding sequences, species discrimination ability is comparatively lower and even insufficient, particularly in some groups (Pochon et al. 2013, 2015, Zhan et al. 2013, Brown et al. 2016, Borrell et al. 2017). For example, in previous studies targeting NIS with 18S, no differentiation has been found among invasive species of molluscs (Pochon et al. 2013, Fletcher et al. 2017, Couton et al. 2019) or between species of the genus *Botrylloides* (Couton et al. 2019). The 18S marker also lacked resolution for tunicate species surveyed through eDNA metabarcoding in Indonesian islands (Huhn et al. 2020).

On the other hand, as the formal DNA barcode region for animals (Hebert et al. 2003), the 5’ segment of COI has been shown to enable species-level identification in the majority of marine metazoan taxa (Bucklin et al. 2011), even using shorter amplicon segments in metabarcoding approaches (Hollatz et al. 2017, Lobo et al., 2017). The high abundance of mitochondrial DNA copies per cell, also increases the success of detecting rare species (Galtier et al. 2009). Nevertheless, particular COI primers have also been reported not to amplify during PCR in some groups due to the lack of conserved primer binding sites (e.g. Copepoda, Cladocera, Nematoda, Creer et al. 2010, Tang et al. 2012, Zhan et al. 2013). Primers targeting the 3’ end of the COI barcode region have been mostly used (Fig. 4), in particular the primer mICOIintF in combination with jgHCO2198 and also, but to a less extent, with dgHCO2198. Both primer combinations have been previously found to perform better across metazoan phylogenetic diversity than the mICOIintF reverse complement with LCO1490 (and its degenerate versions dgLCO1490 and jgLCO1490) (Leray et al. 2013). A primer pair targeting exactly this same region, but using instead LoboR1 (Lobo et al. 2013, 2017) has been shown to produce similar or higher taxa recovery (e.g. Haenel et al. 2017, Ip et al. 2019, Chang et al. 2020), but it has not been tested yet in NIS-focused studies.

At least 12 studies have employed both markers (Table S2) (Borrell et al. 2017, Grey et al. 2018, Koziol et al. 2018, Stefanni et al. 2018, von Ammon et al. 2018b, Couton et al. 2019, Holman et al. 2019, Leduc et al. 2019, Rey et al. 2019, 2020, Wood et al. 2019, Huhn et al. 2020, Westfall et al. 2020); the 18S for detecting a broader range of taxa and the COI for improving metazoan species-level discrimination. In addition, many species have reference sequences for only one of the markers, and thus both markers can act complimentarily (Borrell et al. 2017, Grey et al. 2018, Couton et al. 2019). A few authors also opted for the use of more than one primer pair targeting the same marker to circumvent primer biases and improve the breadth of taxonomic recovery (Pagenkopp Lohan et al. 2016, Lacoursière-Roussel et al. 2018, Leduc et al. 2019).

Only four studies employed other markers than COI or 18S for detecting specific taxonomic groups (rbcL: Zaiko et al. 2015b; 16S: Koziol et al. 2018, Huhn et al. 2020 and Westfall et al. 2020, ITS2; Huhn et al. 2020 and ND4: Westfall et al. 2020) (Table S2). For example, Zaiko et al. (2015b) targeted invasive diatom species using the rbcL marker to survey the ballast water of a vessel crossing the Atlantic Ocean. Without its use, diatoms and yellow-green algae, would remain undetected or highly underestimated (Zaiko et al. 2015b). Koziol et al. (2018) targeted crustacean species by employing specific primers for the 16S mitochondrial gene. This gene remains less explored in bioinvasion studies, but it can provide a good balance between taxonomic resolution and successful amplification breadth for NIS assessments (Westfall et al. 2020). However, the reference library for 16S is more incipient than other widely used markers, such as COI or 18S.

When targeting a broad taxonomic range, the inadequate choice of primers can lead to the underrepresentation or even failed detections of NIS, resulting in false negatives of potential taxa of concern (Darling et al. 2018, von Ammon et al. 2018b, Wood et al. 2019). For instance, DNA metabarcoding employing 18S and COI, failed to detect several bryozoan taxa in a New Zealand’s marina, despite several entries can been found for those species and markers on GenBank (von Ammon et al. 2018b). In addition, in a study in a New Zealand harbour, DNA metabarcoding-based detection rates were low for the polychaete *Sabella spallanzanii*, although this species was highly abundant in the harbour and well represented by reference sequences in genetic databases (von Ammon et al. 2019, Wood et al. 2019). These studies suggest that amplification may have failed due to mismatching primer binding sites or low primer affinity. Possibly the metabarcoding primers differed from the primers used to generate the full-length reference sequences. Thus, the characterization of diverse eukaryotic NIS communities will greatly benefit with the application of multiple marker approaches (Borrell et al. 2018, Grey et al. 2018, Stefanni et al. 2018, von Ammon et al. 2018b, Westfall et al. 2020), multiple primer pairs targeting the same genetic regions (Pagenkopp Lohan et al. 2016, Lacoursière-Roussel et al. 2018) or by designing NIS-specific primers (Ardura et al. 2017, Westfall et al. 2020).

Reliance uniquely on “universal” primers may be insufficient to distinguish between NIS and closely-related native congeners (Westfall et al. 2020). For instance, using 18S, Shaw et al. (2019) were able to identify 19 OTUs known to contain strains of harmful bloom-causing algae in sediments, but could not distinguish between harmful and nonharmful strains within the *Alexandrium* genus. To circumvent this limitation, Westfall et al. (2020) developed a biosurveillance tool tailored for detection of NIS in the Northwest Pacific. These authors designed 8 primer sets targeting multiple gene regions (16S, 18S, COI and ND4) of 4 main groups comprising NIS: Arthropoda, Bryozoa, Mollusca and Ascidiacea. After *in silico* evaluation and refinement, the primers were successfully tested *in situ* with DNA from reference specimens and validated in the field through metabarcoding of water’s eDNA and zooplankton. By combining this approach with a comprehensive sampling of different substrates, they were able to detect 12 NIS in Departure Bay, Pacific Canada, and 7 potential new records, across a wide taxonomic range. However, high intraspecific variation due to multiple sympatric lineages of NIS may lead to failed amplification with species-specific primers, but there is little to no empirical work to support this (von Ammon et al. 2019). Thus, in the future, the use of metabarcoding in biosurveillance may be greatly improved by optimizing the tool for a particular biogeographic region and using a multi-marker approach, which may accelerate its implementation in regular monitoring of NIS worldwide (Westfall et al. 2020).

### 3.7. Sequencing platform and sequencing depth

Three main platforms have been used in studies of NIS surveillance in coastal and marine ecosystems through DNA metabarcoding: 454 Roche (11 publications), Ion Torrent (6 publications) and Illumina MiSeq (27 publications), which has been by far the most widely used (Table S2).

The discrepancy that can be found between sequencing platforms is one of the problems that must be circumvented for a generalized use of DNA metabarcoding data for routine monitoring of biological invasions (Ardura et al. 2015, Zaiko et al. 2015a). In Ion Torrent and 454 Roche, DNA fragments are sequenced-by-synthesis after clonally amplified by emulsion PCR on the surface of microbeads (Shokralla et al. 2012, Zaiko et al. 2015a). Illumina has also adopted a sequencing-by-synthesis approach, but uses fluorescently labelled reversible-terminator nucleotides on clonally amplified DNA templates immobilized on the surface of a flow cell (Shokralla et al. 2012, Zaiko et al. 2015a).

Indeed, in comparative studies, no consistency was found between 454 Roche and Ion Torrent platforms (Zaiko et al. 2015a). Future research should explore further how the use of different platforms may affect the detection of NIS (Deiner et al. 2018, Pagenkopp Lohan et al. 2019, Zinger et al. 2019), especially as DNA metabarcoding gradually becomes a routine surveillance and monitoring tool (Leese et al. 2018). On the other hand, the Illumina’s newest high-capacity platform NovaSeq, which has a sequencing depth 700 times greater than MiSeq, may be more appropriate for the early detection of NIS (Singer et al. 2019). In a recent study, the NovaSeq platform was able to detect many more taxa, even when the sequencing depth matched with that of Illumina MiSeq (Singer et al. 2019). However, the NovaSeq is ca. ten times more expensive than a MiSeq instrument, as well as the sequencing runs cost, and may be out of reach for most laboratories (Singer et al. 2019).

The sequencing depth employed in DNA metabarcoding, although not specified in most studies, can also greatly compromise the results obtained, which may vary depending on the purpose of the study (Grey et al. 2018). Higher sequencing depths may be needed for highly diverse sites or for detecting rare taxa, while lower sequencing depths may be enough for surveying less diverse sites or when using more specific primers. For instance, Singer et al. (2019) found that a typical sequencing depth of 60,000 reads captured about half of the diversity detectable by Illumina MiSeq, in water samples from coastal ecosystems in Canada. For the detection of rare or low abundance species, particularly if a NIS did not spread yet into the ecosystem, a higher sequencing depth may be needed to increase the chance of its early detection. Grey et al. (2018) found that a sequencing depth of 150,000 reads was needed to get ca. 80% of estimated richness, including NIS detection in most water samples collected from ports from 4 different geographic regions (Australia, Canada, Singapore and USA). Whether or not this has a significant impact on a study will depend on the nature of that study (Singer et al. 2019). An evaluation beforehand by sequencing a few samples may help to determine the best sequencing depth to employ in a particular study. But surveys that have an interest in the early detection of NIS could be significantly impacted by the platform and sequencing depth employed.

### 3.8. Bioinformatics pipelines

The bioinformatics workflow is a complex process and highly sensitive to sequence processing parameterization and to taxonomic assignment procedures (e.g. reference databases selection for sequences comparison) (Zhan et al. 2014, Brown et al. 2015, Flynn et al. 2015). In our review, we found that different pipelines have been employed in the bioinformatics workflows (Hatzenbuhler et al. 2017, Scott et al. 2018, von Ammon et al. 2018b) (Table S5), which further complicates comparison among studies.

Most pipelines implemented sequence clustering prior to taxonomic assignments, to account for intraspecific variation and artefactual sequences, produced during PCR and sequencing (e.g. Brown et al. 2016, Ghabooli et al. 2016, Pagenkopp Lohan et al. 2016, 2019, Deiner et al. 2018, Holman et al. 2019, Shang et al. 2019). The algorithms used for OTU clustering can lead to both under- and overestimations of biodiversity, due to inherent challenges in properly aligning and clustering of sequences of variable length (e.g. 18S rRNA gene), and in finding adequate clustering thresholds that broadly reflect the patterns of sequence variability within and among species in phylogenetically diverse communities (Brown et al. 2015, Flynn et al. 2015). A great proportion of the studies employing metabarcoding in NIS surveillance used the UPARSE pipeline for OTU clustering (Table S5) (Brown et al. 2016, Ghabooli et al. 2016, Pagenkopp Lohan et al. 2016, 2019, Deiner et al. 2018, Holman et al. 2019, Shang et al. 2019), which is allegedly among the most reliable methods available for this analysis (Edgar 2013). It is estimated that with UPARSE, OTUs are produced with ≤1% incorrect bases *versus* >3% generated by other methods (e.g. Mothur, QIIME), which may overestimate OTU number (Edgar 2013, Flynn et al. 2015, Ghabooli et al. 2016).

The similarity thresholds used to cluster the sequences can also strongly influence the final output (von Ammon et al. 2018b). Although a 97% threshold has been commonly employed in most biodiversity assessments, and was indeed used in most of the publications analysed in the current review (Table S5) (Zaiko et al. 2016, Pagenkopp Lohan et al. 2017, Pochon et al. 2017), a higher threshold (e.g. 99%) may increase rare taxa detection, which may be better suited to target NIS at early incursion stages (Abad et al. 2016, von Ammon et al. 2018b). However, similarity thresholds can be strongly dependent on the marker and the length of the fragments targeted. While for the 18S rRNA gene (more conserved) an increase from 97% to 99% OTUs clustering may allow species to be split, for typical COI fragments, which display high sequence variability, this alteration may not produce any meaningful increase in the number of species detected (Scott et al. 2018). The same applies with the length of the fragments under analysis, with shorter fragments being probably more sensitive to similarity thresholds than longer fragments.

Scott et al. (2018) explored how sequence processing parameters may influence the taxonomic assignment of 18S sequences from bulk zooplankton samples and the detection of NIS using the 454 platform. By testing 1,050 parameter combinations, the authors found that trimming and sequence quality filtering can strongly affect the final output, and that sequences, particularly at low abundance, can be wrongly classified as noise during denoising, resulting in false-negative errors (Scott et al. 2018). Skipping denoising and clustering proved to be the best option to process metabarcoding sequences for the early detection of NIS, as well as using relaxed filtering. On the other hand, singleton removal had little negative impact in NIS detection, but reduced significantly redundancy and false-positive errors (Scott et al. 2018).

Some other parameters used in the pipeline, such as the minimum number of reads, may be highly relevant as well (Holman et al. 2019). For instance, the false-negative eDNA detection of *Bugula neritina* in water and sediments in UK marinas (but detected through morphology) was found to be a result of the minimum number of reads used in the pipeline, which was not attained for this species (Holman et al. 2019).

Recent bioinformatics algorithms, which use exact sequence variants (ESV) or unique sequences instead of clustering sequences, are becoming preferred approaches to improve the accuracy and sensitivity of taxonomic assignments compared to methods involving pre-clustering of the sequences (Brown et al. 2016, Chain et al. 2016). Brown et al. (2016) detected 6 NIS when reads were BLASTed against their local database, but that remained unnoticed when OTUs were used. The reads matching these NIS formed OTUs with closely related species (low interspecific sequence divergence). Thus, the direct taxonomic assignment of each read without pre-clustering, may be better suited to distinguish between closely related species and more sensitive to detect low abundance taxa, which are crucial elements for detecting early stages of invasive species.

### 3.9. Incompleteness of reference sequences databases for NIS

Another issue that can have a tremendous impact on the success of DNA metabarcoding on NIS surveillance is the availability, taxonomic breadth and reliability of reference sequences databases used for taxonomic assignment (Zaiko et al. 2015c, 2016, Stefanni et al. 2018, von Ammon et al. 2018b, McGee et al. 2019, Rey et al. 2019, 2020). To date, all reference sequences databases remain incomplete or are often tailored for specific genetic markers or taxonomic groups (e.g. PR2 database for the 18S, BOLD database for COI) (von Ammon et al. 2018b, Rey et al. 2019, 2020). GenBank may contain reference sequences from many different genetic markers and includes all domains of life (Briski et al. 2016, Ardura 2019), but it is more prone to errors, since it contains a high number of non-curated data entries (López-Escardó et al. 2018). For example, von Ammon et al. (2018b) were able to detect the invasive species *Sabella spallanzanii* in a New Zealand’s marina by using 2 approaches: 18S with taxonomic assignment to PR2 and COI with taxonomic assignment to BOLD, but the taxonomic assignment in GenBank delivered matches only at the genus level. Couton et al. (2019) were able to detect 3 NIS (*Crepipatella dilatata*, *Watersipora subtorquata* and *Botrylloides diegensis*), when using a custom-designed database, but that were missing when the taxonomic assignments were conducted on GenBank. As long as genetic databases are not complete, the use of different reference databases is probably the best approach for taxonomic assignments (von Ammon et al. 2018b).

The under-representation of some taxonomic groups in reference sequences databases has been the most pointed reason for the failure of NIS detection through DNA metabarcoding (Pochon et al. 2015, Briski et al. 2016, Chain et al. 2016, Ghabooli et al. 2016, Zaiko et al. 2016, Fletcher et al. 2017, Lacoursière-Roussel et al. 2018, von Ammon et al. 2018b, Ardura 2019). For instance, DNA metabarcoding provided similar spatial and temporal trends for zooplankton analysed through morphology, surveyed in the estuary of Bilbao (Spain) (Abad et al. 2016). However, for phytoplankton a different trend was found, probably due to the lack of representative sequences in genetic databases for this group (Abad et al. 2016). Also, the absence of 5 zooplankton species from the DNA metabarcoding results, but detected through morphology in the Baltic sea (Zaiko et al. 2015c), or the substantial number of OTUs matching unknown eukaryotes in biofilms scraped from a New Zealand’s marina (ca. 25%, Pochon et al. 2015), were mostly attributed to the lack of reference sequences in BOLD. In fact, nematodes and tunicates (Chain et al. 2016), Rotifera and Porifera (Lacoursière-Roussel et al. 2018) and bryozoans (Leduc et al. 2019), have been identified as taxonomic groups consistently missing reference sequences in genetic databases, therefore they are more likely to remain undetected when using only the metabarcoding approach.

From the total number of species (n=1083) included in the AquaNIS database (Olenin et al. 2014), which constitutes a worldwide information system on aquatic non-indigenous and cryptogenic species, only ca. 43% and 34% were covered by sequences, in BOLD and PR2 databases, respectively (Rey et al. 2020). Very recently, Duarte et al. (2020), found that from the total number of species included in the AquaNIS and EASIN databases, occurring in European coastal regions, approximately 65% and 55%, respectively, had at least one of the searched barcode markers (COI, 18S, rbcL and matK), in BOLD or GenBank. However, the values of missing barcodes clearly differed among taxonomic groups (i.e. Animalia, Chromista and Plantae) and the barcode markers searched (Duarte et al. 2020). Definitely, expanding genetic databases with reference sequences is mandatory to make the most of the potential of the DNA metabarcoding approach in NIS surveillance in coastal ecosystems. Ideally, this should cover the full sweep of species in the target ecosystem, with a balanced representation of specimens across each species distribution range (native and recipient locations), to account for possible regional variability in targeted barcode genes. As long as reference databases are far from completion, taxonomic assignment of high throughput sequences may be improved by generating local reference sequences libraries (Abad et al. 2016). Abad et al. (2016) was able to increase the taxonomic assignment success from 23.7 to 50.5%, by generating DNA barcodes for local plankton species in the estuary of Bilbao (Spain).

Although completing the gaps in reference libraries is essential, careful compilation, verification and annotation of available sequences is fundamental to assemble large curated and reliable reference libraries that provide support for rigorous species identifications (Couton et al. 2019, Weigand et al. 2019, Leite et al. 2020). To this end, taxonomic expertise remains crucial for an accurate identification of the specimens used to originate the reference sequences. For instance, sequences from *Acartia tonsa*, an invasive species recorded in many ecoregions of the world, formed several distinct clades, some of which clustered with *A. hudsonica* (Lacoursière-Roussel et al. 2018). Viard et al. (2019) also found sequences of *Botrylloides diegensis* erroneously assigned to *B. leachii*. *Ciona intestinalis* and *C. robusta* were recently reclassified and, thus, there are many sequences of *C. robusta* on GenBank falsely attributed to *C. intestinalis* (Gissi et al. 2017, Couton et al. 2019). In addition, species complexes have been uncovered for many invasive species (e.g. *Bugula neritina*, *Watersipora* sp., Mackie et al. 2012, Fehlauer-Ale et al. 2014), and obtaining reference sequences for local specimens can be extremely important to ensure correct species assignments (Couton et al. 2019). These observations have major implications as introduced species can be misidentified as putative native species. Unfortunately, these database errors can be frequent and, thus, auditing existing records and building/compiling curated reference libraries with high-quality sequences (Leite et al. 2020, Fontes et al. 2020), can be just as important as generating new reference sequences for NIS.

## 4. Final considerations

During our review we found several strengths and opportunities of the metabarcoding approach, highlighting its great potential to become a valuable monitoring tool of biological invasions in marine and coastal ecosystems (Fig. 5). These include:

i. improved resolution, enabling the identification of morphologically similar species and eventually cryptic species, due to a higher taxonomic accuracy;
ii. increased ability to detect smaller organisms, undifferentiated life stages and rare species, enabling the detection of potential NIS at early stages of incursion, which are hardly noticeable through morphology;
iii. a higher throughput than morphology-based approaches, allowing the simultaneous analysis of a larger number of samples with a higher spatial and temporal frequency, which is decisive for upscaling survey efforts for early detection of NIS populations in biosurveillance programmes;
iv. faster responses, since less time is required to analyse the samples, and thus, more responsive to immediate regulatory and management needs;
v. possibility to use non-destructive surveys by analysing eDNA from environmental samples, which may be a critical feature to employ for certain target organisms, such as fish, and which in the future may be conducted “*in situ*”;
vi. shorter operator learning and training curves, compared with morphology-based identification;
vii. after optimization to best fit the target ecosystem, subsequent employment can be executed at a relatively low cost per sample, particularly when analysing a large number of samples;
viii. more amenable to automation since the data generated is more suitable to audit and less vulnerable to variability in taxonomic expertise among studies and;
ix. increased taxonomic breadth allowing a whole-ecosystem approach, which documents the composition and distribution of NIS across a wide taxonomic range by surveying different substrates and using different target markers (e.g. zooplankton, protists, harmful algae, sessile and benthic fauna).

**Figure 5.**
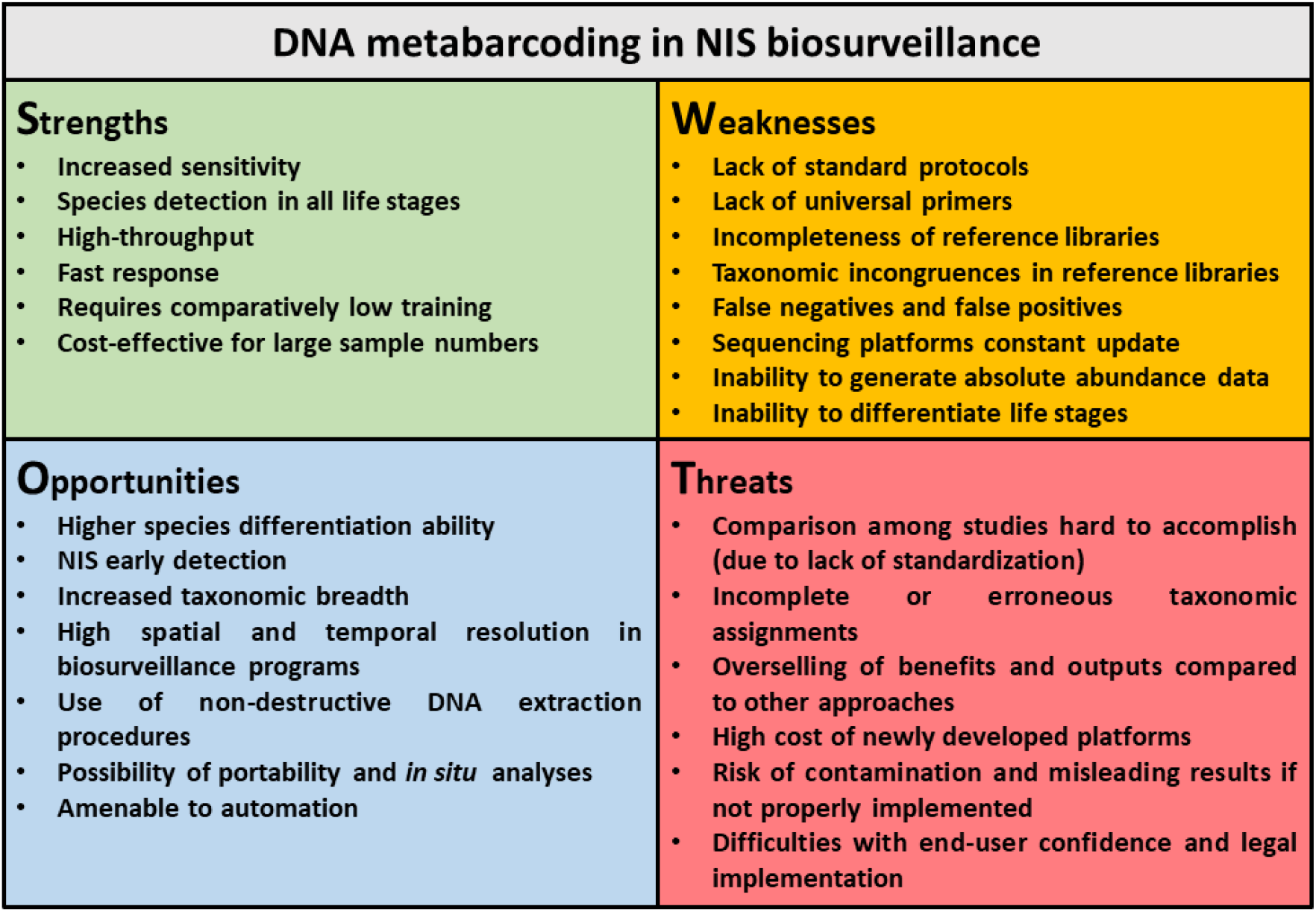
SWOT analysis of DNA metabarcoding in NIS surveillance in marine and coastal ecosystems.

However, some technical limitations still prevail that can be categorized as the main weaknesses and threats, and which must be well acknowledged and managed in order to correct and/or reduce the errors, eventually facilitating a greater adoption of the technique in biosurveillance programs:

i. the workflows of the DNA metabarcoding analytical chain (i.e. sampling design, DNA extraction, target regions and primer sets, high-throughput sequences processing) vary considerably, which makes comparisons across studies hard to accomplish, thus demanding the development of standardized and harmonized approaches;
ii. the lack of effective “universal” primers may compromise species-level detections which are crucial in NIS biosurveillance;
iii. the incompleteness of reference libraries and the existence of taxonomic incongruences, can restrict the diagnosis and detection capacity of NIS at the taxonomic assignment step, leading to false negatives and/or erroneous identifications;
iv. the presence of Type I (false positives) and Type II (false negatives) errors, may lead to data misinterpretations, as both can trigger action or inaction when not required, and, thus, proper controls (negative controls and positive controls) must be designed and employed along the full metabarcoding workflow;
v. the update of sequencing platforms, and high associated running costs, may be out of reach for most laboratories;
vi. the inability to derive absolute abundance figures from DNA-based approaches, and the limited ability to infer relative abundance, among other reasons, due to differences in DNA template copy number among organisms and primer bias and/or preferential amplification of particular species;
vii. despite the fact that a large volume of sequence data is generated through metabarcoding, only a relatively small number of sequences may match potential invaders. Difficulties in determining which species are more likely to establish to prioritize their early eradication or prevent future introductions may arise; and
viii. difficulties with end-user confidence and legal implementation. The advantages of using (e)DNA metabarcoding in NIS surveillance can be obvious for researchers dealing with it, but may seem much more complex and difficult for environmental managers and stakeholders who are new or untrained in DNA-based methods. Thus, it is particularly important to keep the interest and trust of end-users (Darling 2019, Mosher et al. 2020, Sepulveda et al. 2020). For that, it is vital for end-users to be able to: recognize both the power and the limitations of existing tools; to know how (e)DNA metabarcoding works and what are the limitations of the technique and, finally, what can the technique offer to environmental managers beyond what current methodologies already do (Darling 2019). It is also very important that managers know how the tool should be best used, how to interpret the results and how these can influence decisions (i.e. when to act) (Darling 2019, Sepulveda et al. 2020). For that, manuals on best practices and decision-support frameworks, accounting for error minimization and quantification, as well as instruction and training of managers and stakeholders, are imperative (Sepulveda et al. 2020). Clear communication among the diverse partners involved (researchers and managers) in all stages of the research (i.e. protocols definition, samples collection, laboratory analyses, data analyses and results interpretation) is also essential for a successful implementation of molecular detection methods (Mosher et al. 2020).

Until these weaknesses are resolved, NIS metabarcoding is ought to be supported by complementary approaches, where positive detections can trigger in-depth morphological analysis (Borrell et al. 2017) or the application of more targeted molecular methods (e.g. qPCR, ddPCR) (von Ammon et al. 2018b, Wood et al. 2019). Further refinements to the current methodology and improvements in reference sequence databases will facilitate the adoption of this technique for routine NIS surveillance in marine and coastal ecosystems. Continued research is still required to understand more deeply the causes of false positives and false negatives, and minimize their occurrence. Eventually, as methods improve and become standardized, and the cost of sequencing decreases, researchers and natural resources managers will have access to more comprehensive data with much greater resolution and at a fraction of the cost and time of current traditional monitoring surveys.

## Acknowledgements

This work was supported by national funds through the Portuguese Foundation for Science and Technology (FCT, I.P.) in the scope of the project “NIS-DNA: Early detection and monitoring of non-indigenous species (NIS) in coastal ecosystems based on high-throughput sequencing tools” (PTDC/BIA-BMA/29754/2017). We are also grateful to two reviewers for comments and suggestions that greatly improved the manuscript.

