## Supplemental Figures for "Status and prospects of marine NIS detection and monitoring through (e)DNA metabarcoding"

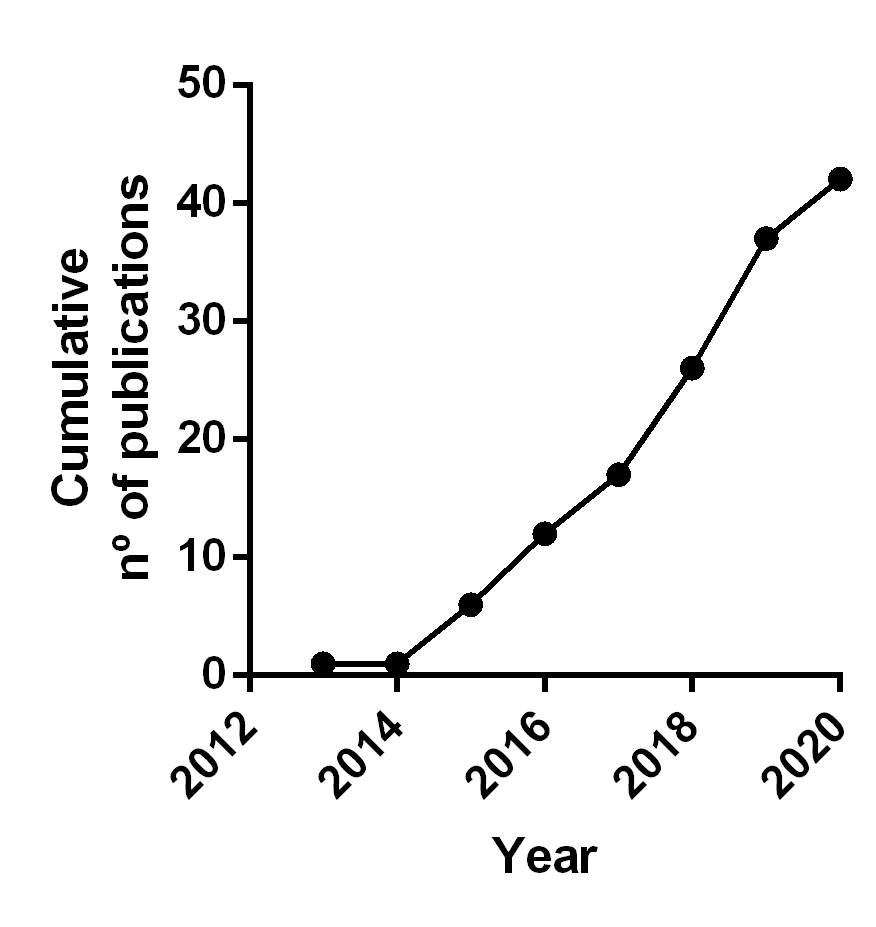

**Figure S1.** Cumulative number of publications dedicated to the use of DNA metabarcoding in NIS surveillance in marine and coastal ecosystems along time. A linear regression was applied to the data indicating an increase of ca. 6.3 publications/year (y=6.33x-12753, r^2^=0.96, p<0.0001).

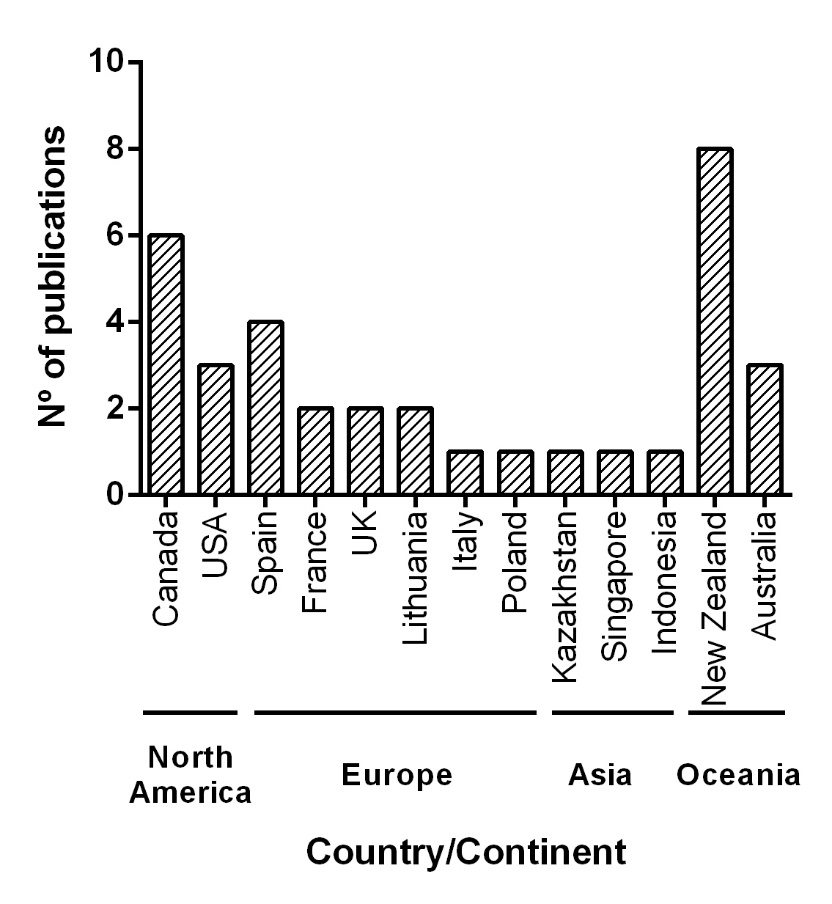

**Figure S2.** Countries where the published studies pertaining the use of DNA metabarcoding were conducted for NIS surveillance in marine and coastal ecosystems.
